# From head to tail: does habitat use drive morphological variation in snakes?

**DOI:** 10.1101/2025.04.25.648294

**Authors:** David Hudry, Anthony Herrel

## Abstract

Convergence is a hallmark of adaptive evolution, yet its extent across different taxa remains unclear. Snakes, with over 4,100 species worldwide, provide a unique model to explore how habitat use influences morphology in a group with an at first sight uniform body plan. We here quantified body, head, and tail shape in over 400 species (∼10% of the global snake diversity) to assess the role of habitat use in shaping morphological variation. Our results reveal significant differences across ecological groups, with habitat use acting as a key driver of phenotypic diversity. Terrestrial species display the highest morphological diversity in contrast to other habitat groups which appear morphologically specialized. For example, whereas arboreal and semi-arboreal species exhibit elongated heads and slender necks, aquatic and semi-aquatic snakes share streamlined bodies and narrow heads. Fossorial and semi-fossorial species, on the other hand, have compact bodies. Surprisingly, morphological convergence remains limited to arboreal, semi-arboreal, and terrestrial habitat groups. Thus, despite strong functional constraints, evolutionary convergence in fossorial and aquatic species is weak, indicating multiple adaptive solutions rather than a single morphological trajectory. Morphological disparity patterns show that non-specialist ecologies generally exhibit greater disparity than highly specialized ones, in accordance with the need of these species to move in different habitats. Our findings underscore the role of ecological constraints in shaping snake morphology and highlight the complexity of adaptation beyond strict convergent evolution.

**Lay summary:** The living environment of a species has been suggested to play a key role in shaping its morphology. Our study aimed to understand whether and how habitat use has influenced the evolution of external body and head shape in snakes. To do so, we gathered a comprehensive dataset of morphological traits from 436 snake species. Our findings reveal that species living in different habitats are morphologically distinct with species living in similar habitats displaying similar morphologies despite the conserved morphology of snakes.

## Introduction

Animals need to adapt to survive and reproduce. The environment in which an animal lives exerts selective pressures on the performance and behavior of an animal, driving selection on underlying morphological traits (Arnold, 1983). Thus, traits that promote escape behavior, feeding efficiency, and reproduction may promote individual fitness and be under strong natural selection. Often, animals subjected to the similar selective pressures will exhibit convergent phenotypes as morphological solutions to maximize performance in a given environment may be limited (Herrel et al., 2008; Deepak et al., 2023). For example, sea snakes hunting burrowing eels show similar body shapes (Sherratt et al., 2018) and aquatic snakes more generally have evolved similar head morphologies minimizing drag in response to hydrodynamic constraints (Herrel et al., 2008; Segall et al., 2016). Such examples of convergent evolution do not only highlight how species adapt to environmental pressures but also provide valuable insights into broader evolutionary mechanisms, such as adaptive evolution, functional trade-offs, and phylogenetic inertia (Speed & Arbuckle, 2017). However, similar phenotypes do not always indicate similar ecological roles as geographical differences in resource availability may exist despite similar morphologies (Grundler & Rabosky, 2014).

Snakes are an excellent model for studying morphological convergence due to their ecological versatility (O’Shea, 2018). They have evolved to thrive in nearly all environments except polar regions. Foraging mode and dietary specialization have been suggested to be the main drivers of the evolutionary success of snakes (Title et al., 2024). This dietary diversity surged after the Cretaceous mass extinction (Grundler & Rabosky, 2021), with ecological opportunism and behavioral flexibility driving adaptive radiations to different diets. For example, distantly related snail-eating snakes, such as those in the genera *Dipsas*, *Sibon*, and *Pareas*, have converged on similar skull shapes (Pandelis et al., 2023). Similarly, habitat-driven head shape convergence occurs in *Boidae* and *Pythonidae* (Esquerré & Keogh, 2016), and selective pressures associated with foraging drive morphological evolution in sea snakes (Sherratt et al., 2018) and homalopsid snakes (Fabre et al., 2016). Furthermore, snakes exhibit different activity patterns, with species being diurnal, nocturnal, or crepuscular, driven by prey availability and thermoregulatory needs (Huey & Kingsolver, 1989; Lettoof et al., 2023). This array of adaptations has made snakes one of the most species-rich groups of reptiles, with over 4,000 currently described species (Pyron & Burbrink, 2012).

Previous studies have explored convergence in snakes in relation to habitat use. For example, arboreal snakes tend to have slender bodies and long tails for balance and movement in trees (Lillywhite & Seymour, 2011; Jayne et al., 2015; Alencar et al., 2017), while aquatic snakes have more streamlined heads to reduce drag and capture prey efficiently (Segall, 2019). Fossorial snakes are adapted for burrowing, with reinforced heads, cylindrical bodies, and reduced ventral scales to minimize friction (Strong et al., 2021). Similarly, sand-dwelling snakes show streamlined heads with countersunk jaws, cylindrical bodies, and reduced ventral scales helping them move through sand (Sharpe et al., 2015). Terrestrial snakes, on the other hand, often have elongated bodies and enlarged ventral scales for efficient movement on land (Brischoux & Shine, 2011). Unfortunately, most studies on convergence have focused on specific families of snakes, such as Natricinae (Deepak et al., 2023), Boidae (Esquerré & Keogh, 2016), Dipsadinae (Pandelis et al., 2023), and Hydrophiidae (Sherratt et al., 2018). Yet, whether morphological convergence can be observed across snakes more generally, and to what degree remains largely unexplored.

Our study aims to address this by analyzing a phylogenetically and ecologically diverse yet comprehensive data set composing roughly 10% of the overall species diversity. We predict that snakes occupying similar habitats will exhibit significant morphological convergence irrespective of their phylogenetic affinity. Based on previous studies we predict that arboreal species will have slender bodies and narrow heads, that aquatic snakes will have more robust bodies and heads with dorsally positioned nostrils, and that fossorial species will show adaptations for burrowing with dorsally positioned nostrils and eyes and short cylindrical bodies. Sand-dwelling snakes are expected to show partial convergence with fossorial species, while semi-aquatic, semi-fossorial, and semi-arboreal species will likely exhibit intermediate traits between specialist categories. We also predict that species using multiple habitats will show greater morphological disparity, reflecting the diversity of ecological niches they occupy. Habitat specialists such as fossorial snakes, on the other hand, are expected to show lower morphological disparity due to the strong physical constraints imposed by their locomotor environment.

## Materials and Methods

### Sample and ecological classification

We collected data for a total of 984 specimens of 436 species of snakes (approximatively 2 individuals/species) housed in three European Natural History Museums: The National Natural History Museum of Paris (181 specimens), the Alexander Koenig Museum of Bonn (611 specimens), and the Museum of Natural Sciences of Brussels (192 specimens). All measurements were taken on adult specimens where possible. For some species only young or juvenile individuals were available and for others subadults were measured for practical considerations (e.g. large species of boas and pythons; Table S1). Our sampling aimed to cover the phylogenetic and ecological diversity across snakes while being constrained by the availability of specimens in these collections.

Habitat classifications were defined based on the literature (Spawls et al., 2002; Whitaker & Captain, 2004; Campbell & Lamar, 2004; Marais, 2004; O’Shea, 2005; Vogel, 2006; Glaw & Vences, 2007; Bartlett & Bartlett, 2009; Das, 2010; O’Shea, 2018; Allen, 2019; Chippaux, 2019; Pietersen et al., 2021; Egan, 2022; Das, 2023). We created eight habitat groups that included 178 terrestrial, 9 sand-dwelling, 56 arboreal, 26 aquatic, 16 fossorial, 65 semi-arboreal, 48 semi-aquatic, and 40 semi-fossorial species. These categories were based on the physical demands imposed by the environment on the snake’s body and head (Table S2). For example, the same hydrodynamic constraints apply to snakes living in the ocean or in fresh water, causing us to group these together.

### Morphometric data

Morphometric data were collected by measuring 13 morphological traits (Table S3). The traits were selected based on the literature on morphometric studies in snakes (Hibbitts & Fitzgerald, 2005; França et al., 2008; Pyron & Burbrink, 2009; Grundler & Rabosky, 2014; Burbrink & Myers, 2015; Cavalheri et al., 2015; Esquerré & Keogh, 2016; Reynolds et al., 2016; Manjarrez et al., 2017; Alencar & Quental, 2022). Snout-vent length and total length were measured using a measuring tape (± 1 cm). The remaining traits were measured with a digital caliper (Mitutoyo © 150 mm 500-181-30; ± 0.02 mm). Before starting data collection, a repeatability test was performed with individuals of the species *Trimeresurus albolabris*.

We measured 10 randomly selected individuals ten times and performed a PCA to explore whether the measurement error was high or not. Our PCA results showed that measurement error is low and that each individual occupies a distinct position in the morphospace (Figure S1).

### Phylogenetic comparative analyses

For each snake species, mean morphological values were calculated as comparative analyses are conducted at the species level. A Log10-transformation was applied to render the data normal and homoscedastic. To perform phylogenetic comparative analyses, we used the phylogenetic tree of Title and co-authors (Title et al., 2024) imported as a newick file. The tree was pruned to include only species present in our dataset using the *drop.tip* function from the *ape* package (Paradis & Schliep, 2019). In total, 433 species were included in the morphometric analysis. Some species were missing from the original tree (14 in total) and were replaced by closely related taxa present in the phylogenetic tree of Title et al. (2024; Table S4). Three species that were measured were ultimately not included in the analysis as they were not included in any phylogeny: *Hydraethiops melanogaster*, *Hydraethiops laevis,* and *Hypoptophis wilsoni*.

Phylogenetic generalized least squares (PGLS) regressions were performed to examine the relationship between traits while controlling for phylogeny using the *caper* package (Orme et al., 2013). Residuals from the PGLS models of each trait regressed against snout-vent length were extracted to assess trait variation independent of size. Next, a phylomorphospace was created using the *ggphylomorpho* package (Barr, 2017), integrating the phylogenetic tree with morphological data. A phylogenetic multivariate analysis of co-variance (Phylo-MANCOVA) with snout-vent length as our co-variate was then conducted using the *mvMORPH* package (Clavel et al., 2015). We used a Wilks’ Lambda test with 1,000 simulations. Differences between ecological groups means were subsequently assessed using Tukey’s post-hoc tests.

### Phylogenetic signal

Blomberg’s K statistic (Blomberg et al., 2003) was calculated in R using the *phytools* package (Revell, 2012) and the *geiger* package (Pennell et al., 2014) to evaluate the phylogenetic signal present in the morphological traits. This statistic measures the extent to which traits exhibit a phylogenetic signal, quantifying the degree of trait similarity among closely related species compared to what would be expected under a null model of trait evolution. A K value close to 1 indicates a strong phylogenetic signal, meaning that closely related species have more similar traits than would be expected by chance. Conversely, values significantly less than 1 indicate a weaker phylogenetic signal, suggesting that trait variation is less influenced by phylogenetic relationships.

### Convergence among niches

The occurrence of convergent evolution among species within the same habitat use was examined using the *search.conv* function from the *RRhylo* package (Castiglione et al., 2019). This function identifies convergent evolutionary changes by assessing whether branches associated with similar ecological states exhibit greater phenotypic similarity than expected under a Brownian motion model of evolution. The analysis was performed using a time-calibrated phylogeny, allowing for the evaluation of both the occurrence and timing of convergence.

### Disparity among and between niches

Morphological disparity across ecological niches was assessed to determine whether certain habitat groups exhibit higher or lower phenotypic variability. Disparity was quantified as the sum of variances across a set of morphological traits. To test for differences in morphological disparity among habitat groups, we used the *dispRity* package (Guillerme, 2018). First, species were grouped according to their ecological niches, and bootstrap resampling (1000 iterations) was applied to estimate disparity distributions. Statistical comparisons were conducted using pairwise Wilcoxon tests with Bonferroni correction to account for multiple comparisons. Additionally, disparity was calculated by summing trait variances for each species, allowing for a finer-scale visualization of disparity patterns across ecological niches. These results were visualized using violin plots, where each point represents an individual species, and the distribution of disparity within each niche is displayed

## Results

### Variation in body and head shape

In our principal component analysis (PCA), 90% of the variation in body and head shape is explained by the first two principal components (PC1: 85.11%; PC2: 4.91%; Figure 1). PC1 mainly summarizes the variation in overall head shape (Table S5), with the highest loadings observed for jaw length (0.291), interocular distance (0.285), head height (0.287), and head width (0.284). This indicates that PC1 captures variation in head length, width, and height. A low PC1 value is associated with a relatively shorter, narrower jaw and head, while a high PC1 value corresponds to a relatively longer, wider jaw and head. PC2, on the other hand, captures variation in body proportions (Table S5). The greatest loadings were observed for snout-vent length (-0.537) and tail length (-0.645). This axis further suggests that individuals with a greater body and tail length tend to have narrower heads. Conversely, higher values of head width (0.309) and quadrate length (0.274) are associated with shorter bodies. PC3 explains additional variation linked to specific proportions of the head and body (Table S5). The highest loadings for PC3 are observed for head length (0.265), rostral-ocular distance (0.274), and neck width (-0.441). This indicates that PC3 captures variations in head length and associated structures. A high value of PC3 is associated with a longer head, particularly in the rostral region (i.e. a longer snout), while a negative value corresponds to a shorter snout and a wider neck. Interestingly, most habitat groups occupy a large part of the morphospace, especially terrestrial species that are equally distributed across each axis.

**Figure. 1:**
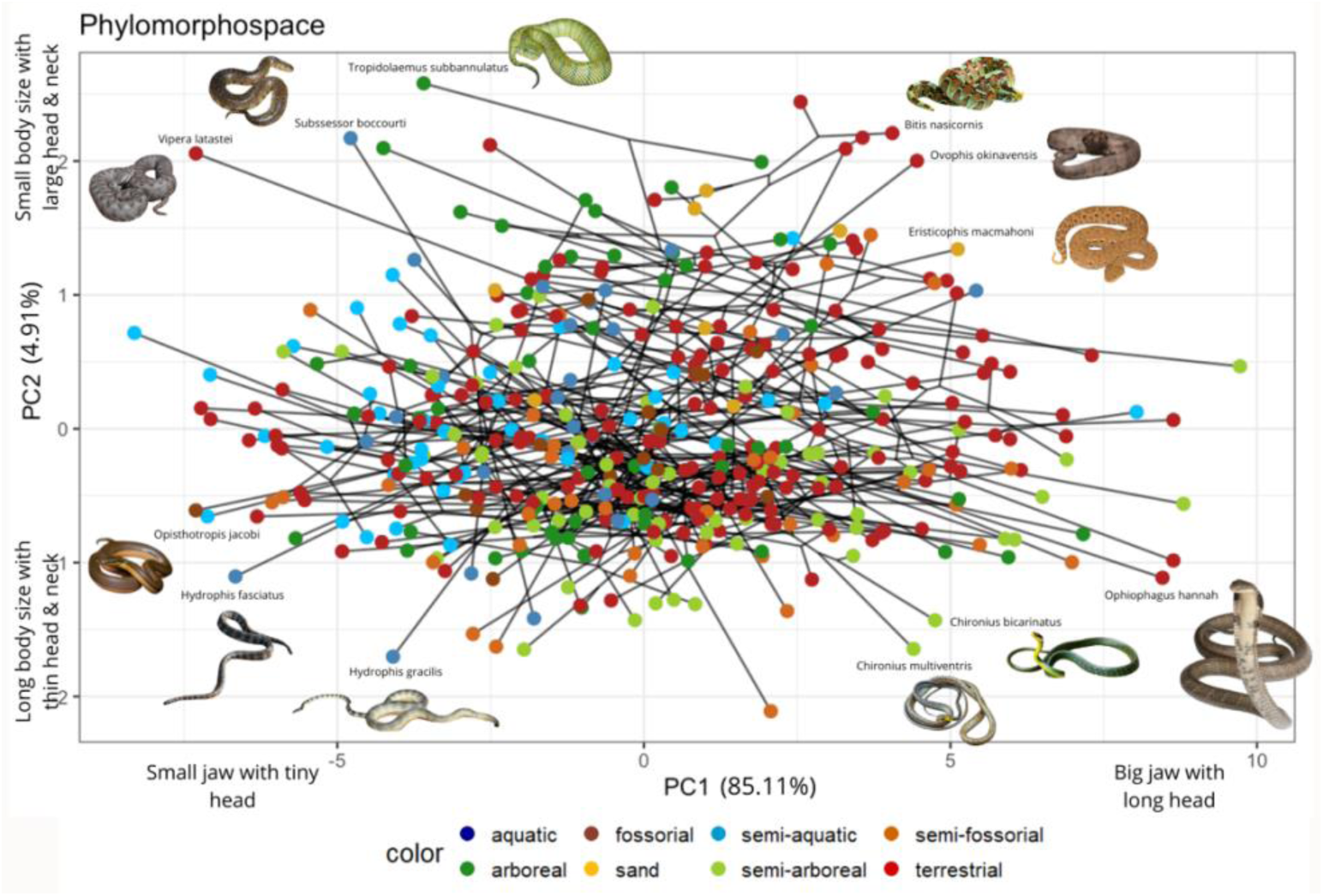
Phylomorphospace illustrating body and head shape variation across 436 species of snakes. The phylogeny is plotted in the morphospace described by the first two axes (PC1 vs. PC2). Nodes are colored based on symmetric rates ancestral-state estimation for habit mapped to the phylogeny.

### Phylogenetic signal

Blomberg’s K test revealed that the phylogenetic signal was low for all traits (Table S6). This indicates that morphological traits among closely related species are less similar than expected and suggests adaptive trait variation.

### Differences between habitat groups

Morphological traits were different between ecological groups when taking into account variation in snout-vent length (Table 1). The effect of snout-vent length and the interaction between snout-vent length and habitat group was also significant suggesting different allometric patterns in species using different habitats (Table 1). Moreover, Tukey post-hoc tests revealed significant size differences between snakes utilizing different habitats (Table 2). Arboreal and semi-arboreal species were significantly longer than all other ecotypes. Semi-aquatic species were also longer than aquatic, semi-fossorial, fossorial, and sand-dwelling snakes. Terrestrial species exhibit medium values in snout-vent length compared to fossorial and sand-dwelling species. Body shape (i.e. dimensions respective of variation in snout-vent length) also differed significantly among habitat groups (Table 2).

**Table 1:** Phylogenetic MANCOVA results, including degrees of freedom (d.f.), Wilks’ Lambda, the F-value, and the P-value.

| Model | df | Wilks' Lambda | *F* | *P* |
| --- | --- | --- | --- | --- |
| Ecological group | 7 | 0.38 | 5.07 | $P < 0.01$ |
| Snout-vent length | 1 | 0.82 | 7.26 | $P < 0.01$ |
| Interaction | 7 | 0.68 | 1.90 | $P < 0.01$ |

**Table 2:**
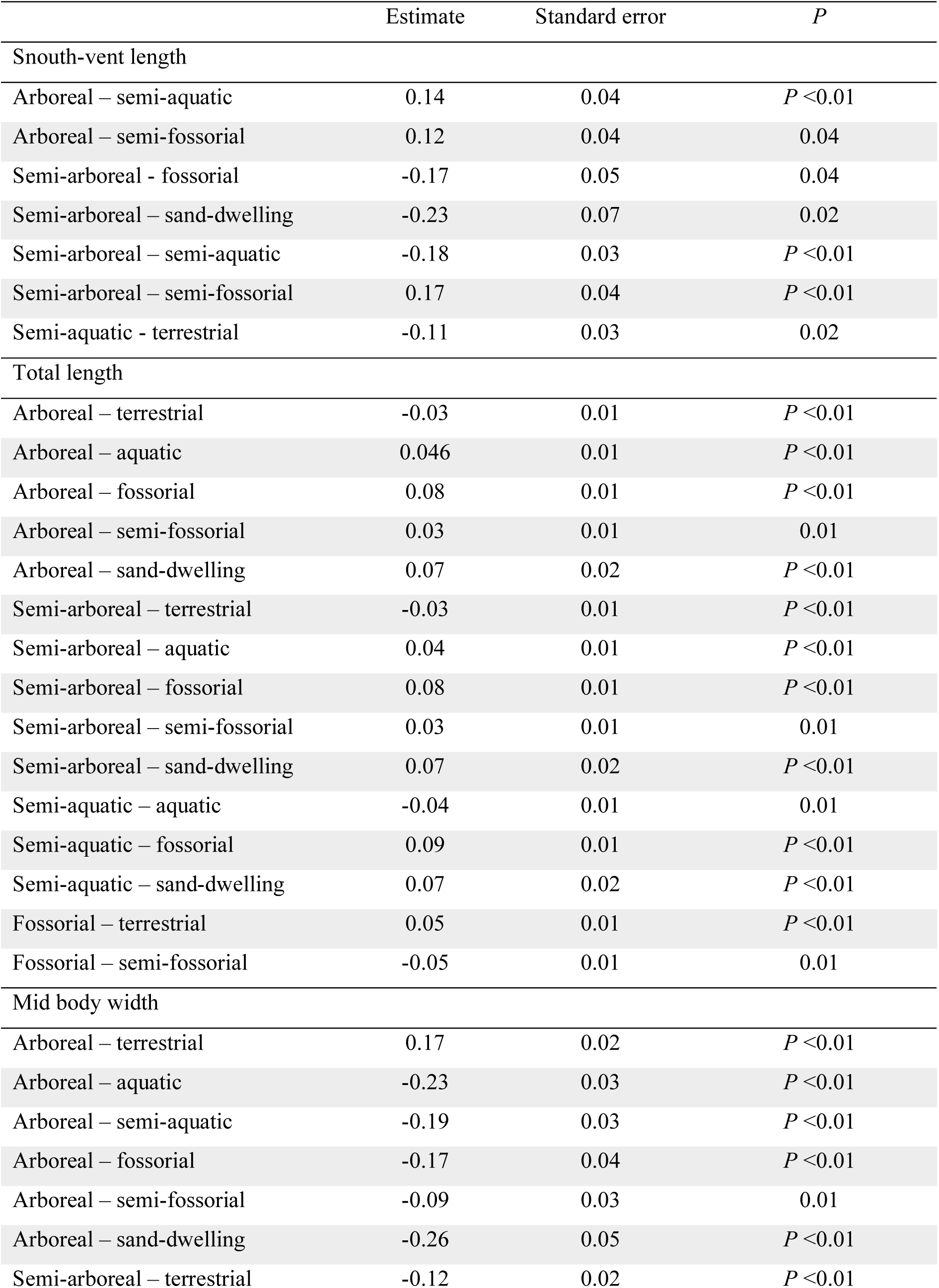
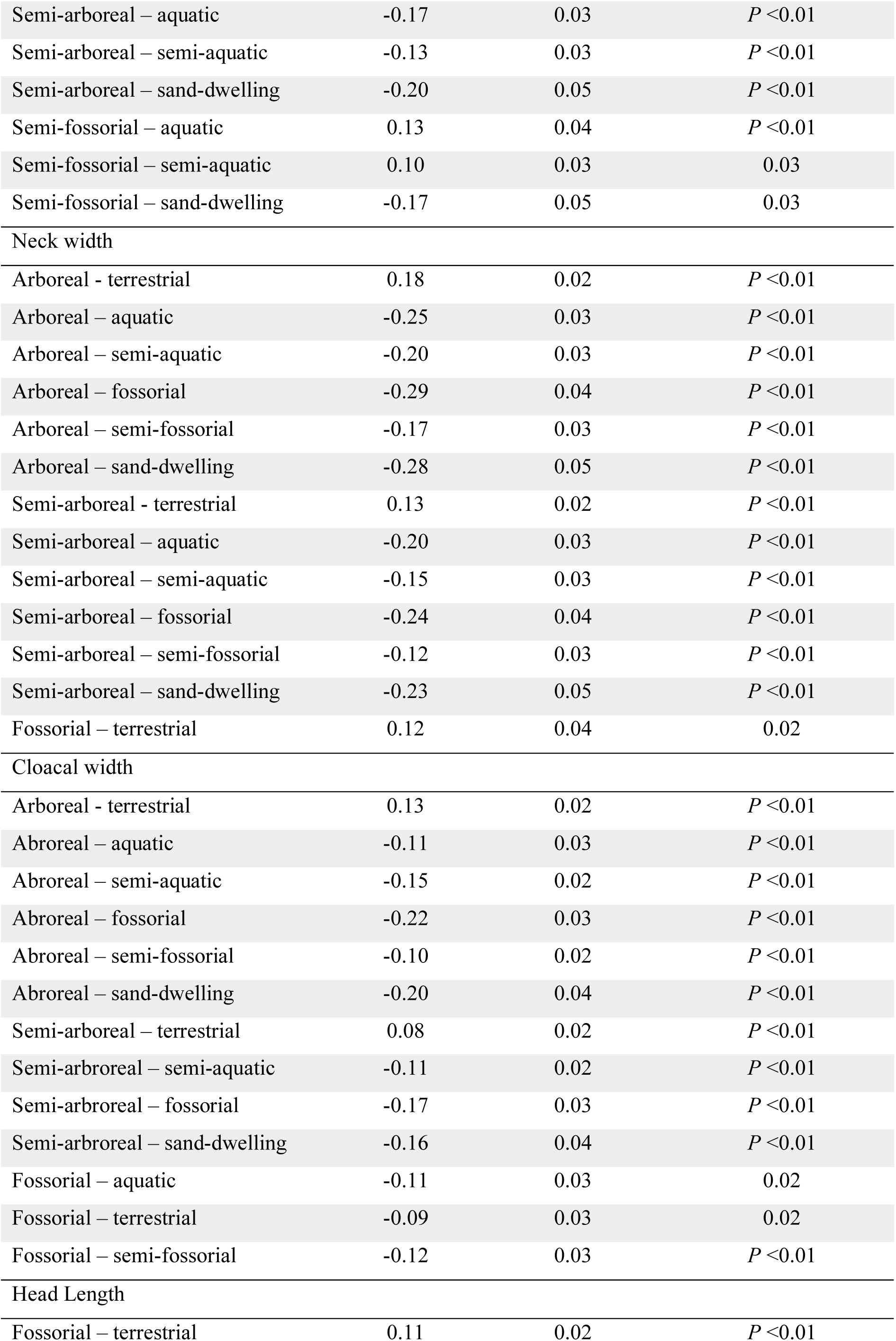
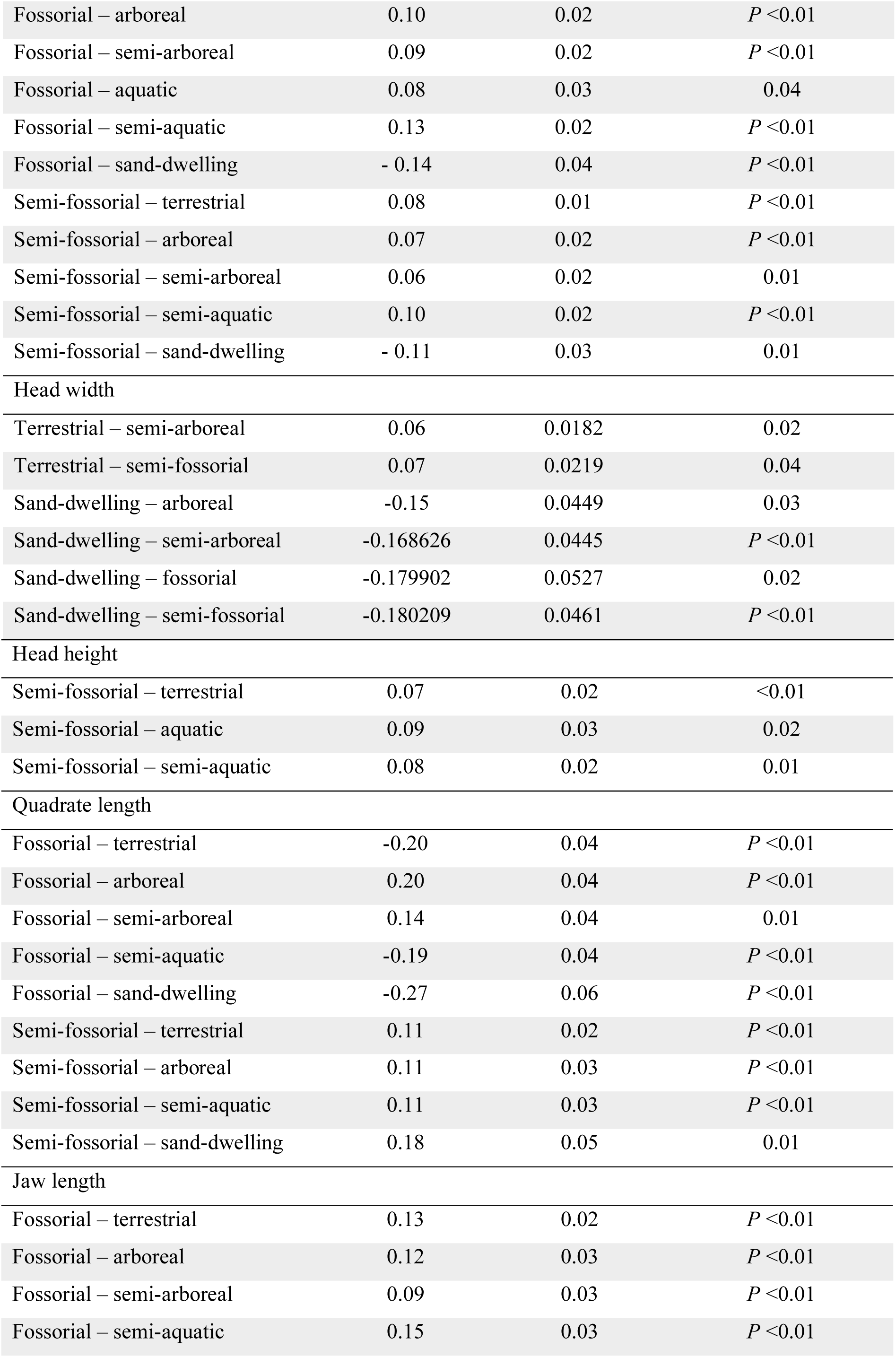
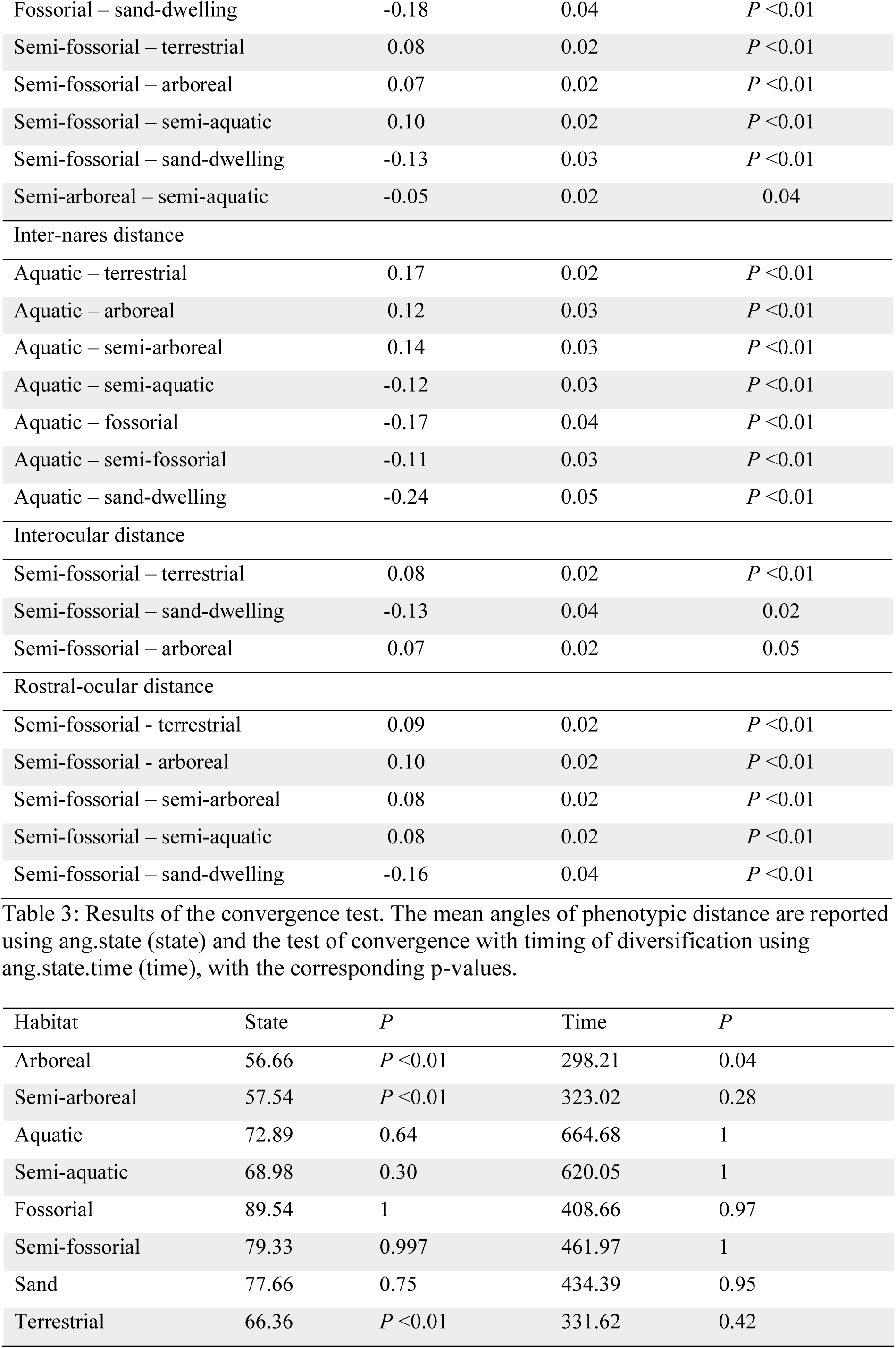
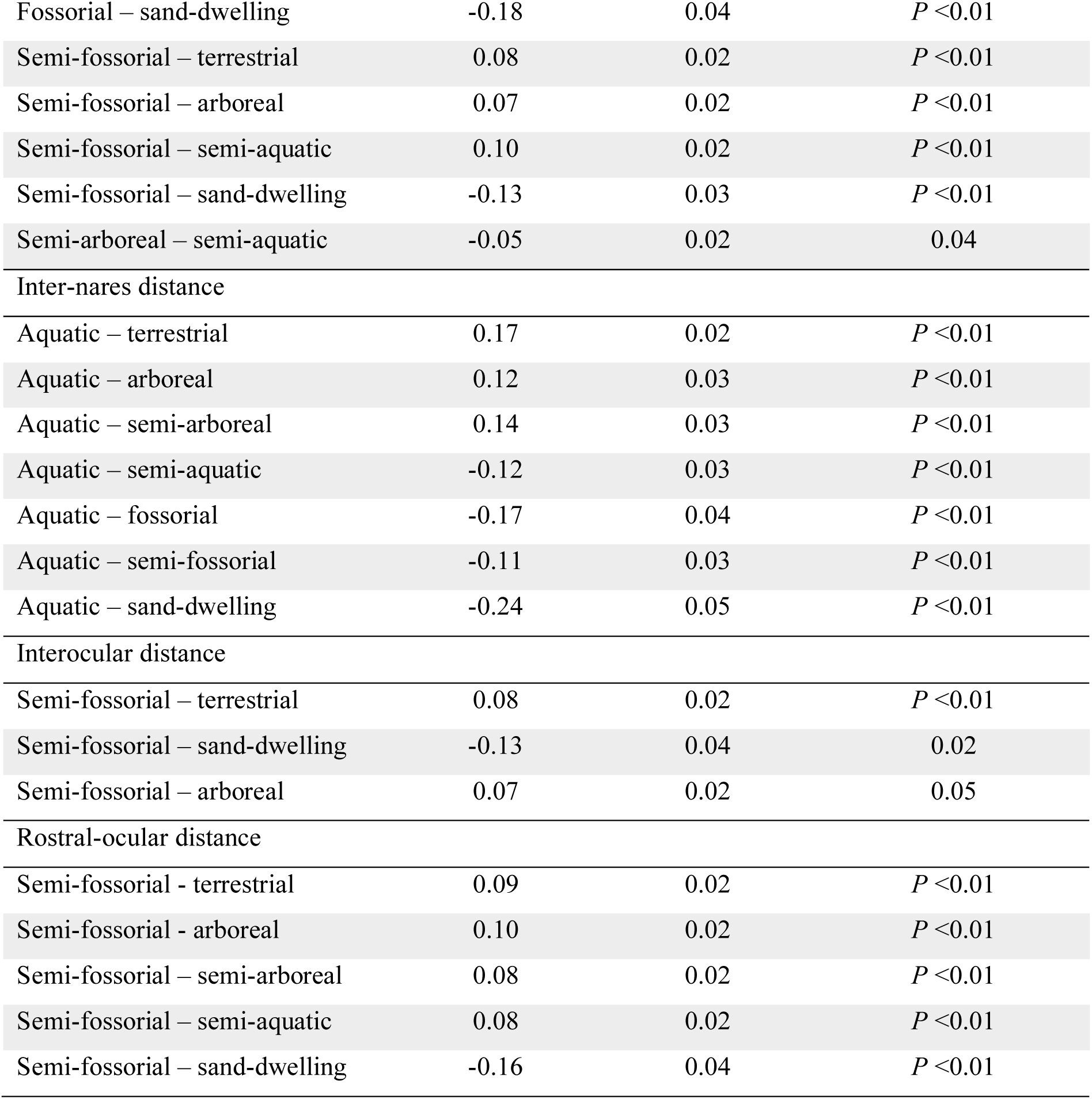
Tukey post-hoc test results, including estimate, standard error, and *P*-values comparing snout-vent length and size-corrected morphological traits and total length measurements across different habitat groups. Only significant comparisons are indicated.

Sand-dwelling and fossorial species had the widest bodies relative to their snout-vent length. Aquatic and semi-aquatic species exhibit the second widest bodies and were different from terrestrial, semi-fossorial, arboreal, and semi-arboreal species. Arboreal and semi-arboreal species were relatively thinner compared to other snakes (Table 2).

Significant differences were also observed in head dimensions (Table 2). Terrestrial species had longer, wider, and taller heads compared to semi-fossorial species and had longer heads than fossorial species. Arboreal and semi-arboreal species also had longer heads than fossorial species, and had longer, wider, and taller heads compared to semi-fossorial species. Moreover, fossorial, and semi-fossorial species had shorter lower jaws and quadrates compared to terrestrial, arboreal, and semi-arboreal species (Table 2). Sand-dwelling species had longer, wider, and taller heads compared to all ecotypes. Semi-aquatic species also had longer heads than fossorial species, and had wider, and taller heads compared to fossorial and semi-fossorial species, respectively. Moreover, aquatic species had a relatively lower inter-nares distance compared to terrestrial, semi-arboreal, and arboreal species. Finally, arboreal species had a relatively greater interocular and eye-nares distance than semi-aquatic and semi-fossorial species.

### Convergence among and between niches

The convergence tests show an overall tendency for species that occupy a given habitat to be convergent (*ang.state*). This was significant for arboreal, semi-arboreal, and terrestrial groups only, however (*P* < 0.05; Table 3). In contrast, for the timing of convergence (*ang.state.time*) the result was significant only for arboreal species (Table 3). This suggests that, in most cases, the evolution of niches was not synchronous, and species converged to their respective morphology at different time points.

**Table 3:**
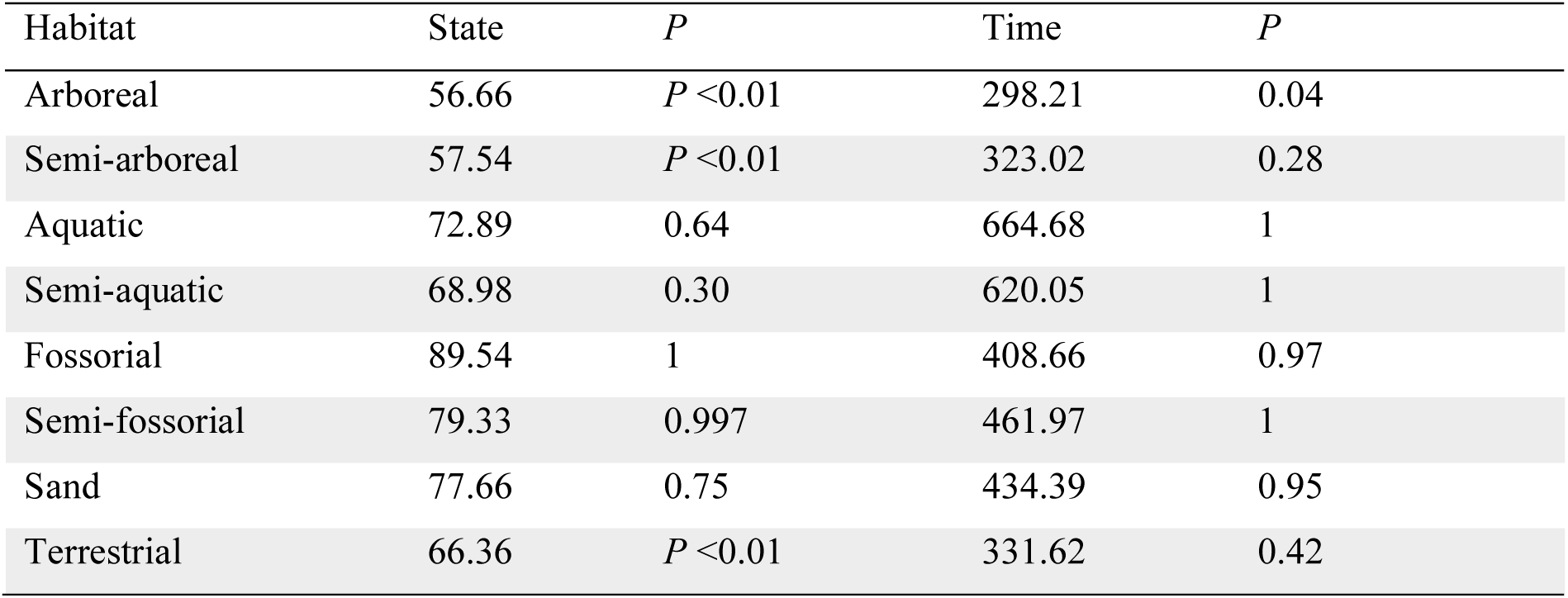
Results of the convergence test. The mean angles of phenotypic distance are reported using ang.state (state) and the test of convergence with timing of diversification using ang.state.time (time), with the corresponding p-values.

### Disparity among and between niches

Semi-fossorial species showed the highest disparity (0.516), followed by terrestrial (0.499) and semi-arboreal (0.448) species (Table 4; Figure 2). The lowest disparity was exhibited by fossorial (0.277) and sand-dwelling (0.291) species. Arboreal, aquatic, and semi-aquatic species exhibit intermediates scores (0.401, 0.351 and 0.386 respectively). Pairwise comparisons indicate significant differences in disparity among nearly all niches (Table 5; Figure 3). Arboreal, semi-arboreal, aquatic, and semi-aquatic habitat groups showed significant differences from the other habitat groups (all *P* < 0.05). Likewise, terrestrial, fossorial, semi-fossorial, and sand-dwelling habitat groups showed significant differences in disparity with two notable exceptions: terrestrial and semi-fossorial groups did not differ in disparity and similarly fossorial and sand-dwelling groups also did not differ.

**Figure. 2:**
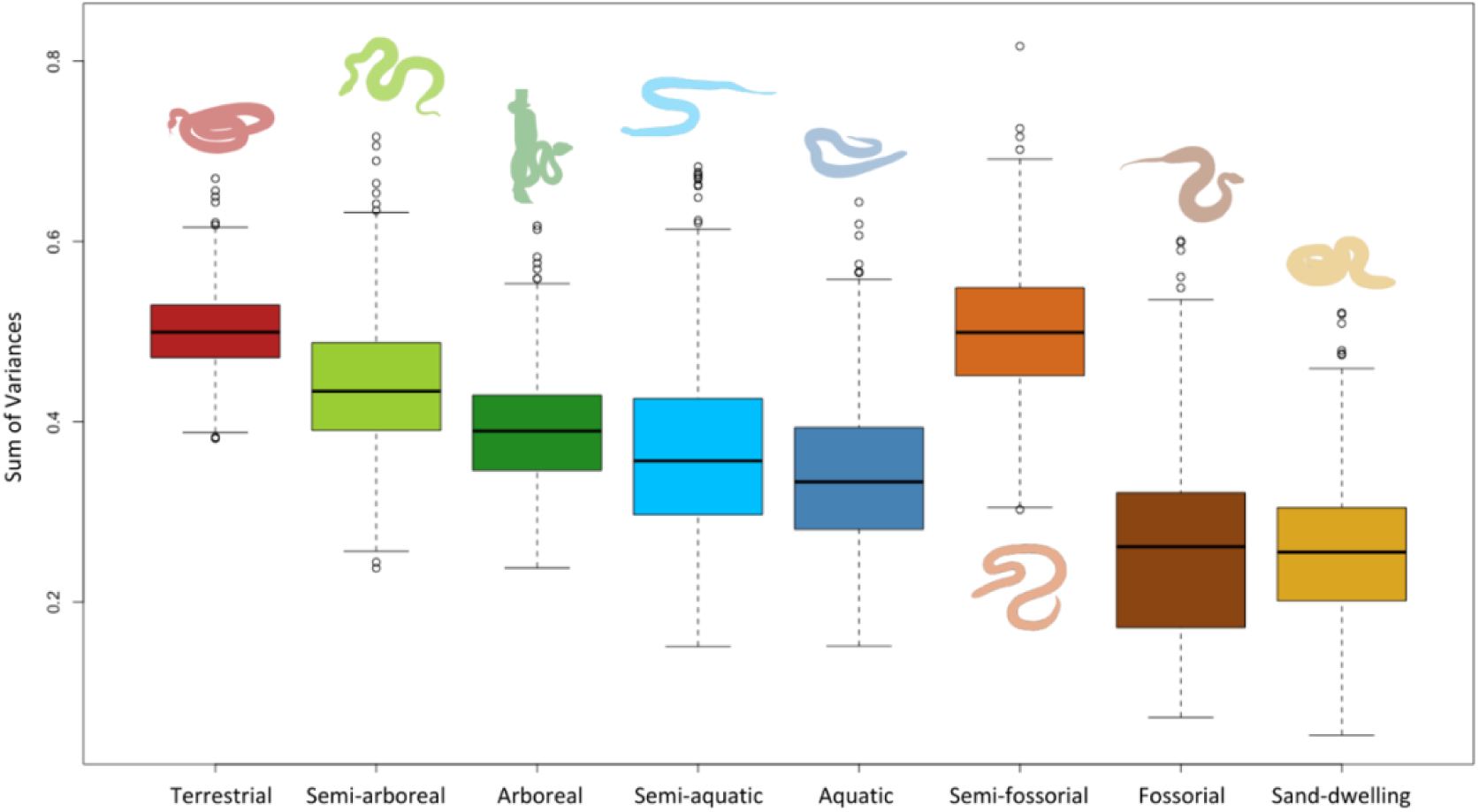
Box plot illustrating the sum of morphological disparity variances across eight habitat groups.

**Figure. 3:**
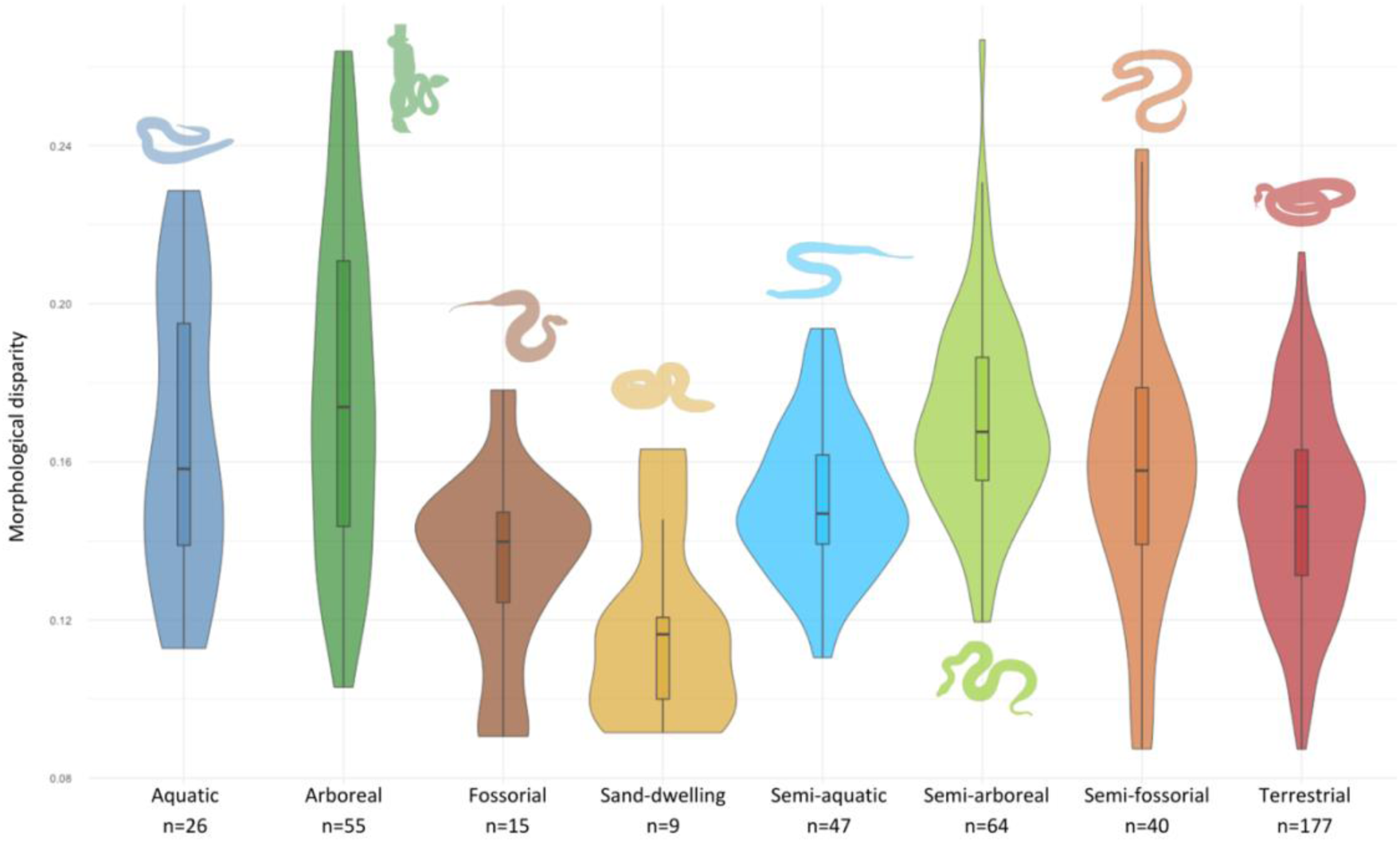
Violin plot illustrating the distribution of morphological disparity for the eight habitat groups

**Table 4:** Results of the morphological disparity using the sum of variances. The table reports the number of species in each habitat group (N), the observed disparity (obs), and the median value from bootstrap resampling (bs.median).

| Habitat | N | obs | bs.median |
| --- | --- | --- | --- |
| Arboreal | 55 | 0.401 | 0.390 |
| Semi-arboreal | 64 | 0.448 | 0.434 |
| Aquatic | 26 | 0.351 | 0.333 |
| Semi-aquatic | 47 | 0.386 | 0.357 |
| Fossorial | 15 | 0.277 | 0.261 |
| Semi-fossorial | 40 | 0.516 | 0.499 |
| Sand | 9 | 0.291 | 0.255 |
| Terrestrial | 177 | 0.499 | 0.499 |

**Table 5:** Results of the pairwise comparison of disparity between habitat groups with Bonferroni correction.

| Niche | *P* |
| --- | --- |
| Terrestrial – semi-arboreal | $P < 0.01$ |
| Terrestrial – arboreal | $P < 0.01$ |
| Terrestrial – semi-aquatic | $P < 0.01$ |
| Terrestrial – aquatic | $P < 0.01$ |
| Terrestrial – fossorial | $P < 0.01$ |
| Terrestrial – semi-fossorial | 0.723 |
| Terrestrial – sand-dwelling | $P < 0.01$ |
| Arboreal – semi-arboreal | $P < 0.01$ |
| Arboreal – aquatic | $P < 0.01$ |
| Arboreal – semi-aquatic | $P < 0.01$ |
| Arboreal – fossorial | $P < 0.01$ |
| Arboreal – semi-fossorial | $P < 0.01$ |
| Arboreal – sand-dwelling | $P < 0.01$ |
| Aquatic – semi-arboreal | $P < 0.01$ |
| Aquatic – semi-aquatic | $P < 0.01$ |
| Aquatic – fossorial | $P < 0.01$ |
| Aquatic – semi-fossorial | $P < 0.01$ |
| Aquatic – sand-dwelling | $P < 0.01$ |
| Fossorial – semi-arboreal | $P < 0.01$ |
| Fossorial – semi-aquatic | $P < 0.01$ |
| Fossorial – semi-fossorial | $P < 0.01$ |
| Fossorial – sand-dwelling | 0.270 |

## Discussion

Terrestrial, arboreal, and semi-arboreal species display a broad distribution across the phylomorphospace. Terrestrial species show extensive morphological variation, as predicted and confirmed by the disparity results, aligning with studies that demonstrate that phylogenetic history and environmental gradients influence morphological diversity in terrestrial snakes (França et al., 2008; Burbrink & Myers, 2015). In contrast, arboreal species, although distributed widely along PC1, cluster around the middle of PC3, suggesting that despite variation in body size and shape, they share common head and neck features such as a thin neck, an elongated head, and widely spaced eyes. This pattern is consistent with studies that found that arboreal vipers, boas, and pythons exhibit constrained morphological evolution, favoring elongated and slender body shapes that enhance maneuverability in three-dimensional environments (Esquerré & Keogh, 2016; Alencar et al., 2017).

Aquatic, semi-aquatic, fossorial, and semi-fossorial species are more tightly clustered in most phylomorphospace plots. Aquatic and semi-aquatic species cluster particularly in the lower left of the PC1 vs. PC3 plot, indicating shared traits such as streamlined bodies, shorter sizes, and narrower heads with wider necks (Figure S2). These findings are likely due to similar hydrodynamic adaptations in aquatic snakes including a streamlined body and dorsally positioned nostrils as observed in previous studies (Brischoux & Shine, 2011; Segall et al., 2016). Fossorial and semi-fossorial species also exhibit tight clustering, especially along the lower end of the PC2 vs. PC3 axes, and show shorter heads, narrower necks, and compact bodies (Figures S3). These results are in accordance with the convergent evolution of compact skulls facilitating burrowing found in other studies (Strong et al., 2021).

Our analysis revealed significant differences in body size across habitat groups (Table 2). Arboreal, semi-arboreal, and semi-aquatic species tend to be significantly longer than aquatic, fossorial, semi-fossorial, and sand-dwelling species. Among them, arboreal and semi-arboreal snakes exhibit the greatest total length, whereas fossorial species have more compact bodies, likely facilitating burrowing and movement through soil. Sand-dwelling species also exhibit smaller bodies, possibly to minimize friction with the substrate. Conversely, aquatic species, are generally smaller in body size. Significant differences in body shape were also detected (Table 2). Sand-dwelling and fossorial species had the widest bodies relative to their length, likely optimizing the space available for the epaxial muscles powering the forward movements during burrowing (Gans, 1970; Newman & Jayne, 2018). Aquatic and semi-aquatic species were relatively wider than terrestrial, semi-fossorial, arboreal, and semi-arboreal species. A wider body presents a greater surface area perpendicular to the movement direction while undulating and may thus imply more efficient swimming. In contrast, arboreal and semi-arboreal species were the most slender, likely enhancing maneuverability and allowing for efficient gap-bridging in complex three-dimensional environments (Jayne, 2020).

Sand-dwelling, terrestrial, arboreal, semi-arboreal and semi-aquatic species had significantly longer heads compared to fossorial and semi-fossorial species (Table 2). Sand-dwelling species also exhibited wider and taller heads than fossorial and semi-fossorial species. Overall, sand-dwelling species displayed the most robust heads, being significantly longer, wider, and taller relative to their snout-vent length. Aquatic and semi-aquatic species had a similar head width and height, with semi-aquatic species displaying relatively taller heads than aquatic snakes. We found similar patterns as Fabre et al. (2016), where aquatic and semi-aquatic species were shown to have comparable head widths. However, semi-aquatic snakes tend to have relatively taller heads, possibly reflecting adaptations to their dual-environment lifestyle. In addition, aquatic species had the shortest inter-nares distance among all groups, likely due to the dorsal placement of their nostrils, which allows breathing while the body remains submerged, an adaptation to aquatic lifestyle (Silva et al., 2017). Similarly, semi-aquatic species had relatively narrow heads, resembling their fully aquatic counterparts. Fossorial and semi-fossorial species had the shortest jaws and quadrates, likely reflecting adaptations for substrate penetration and reducing the cost of burrowing (Table 2). Semi-fossorial species exhibited the shortest interocular distances among all groups suggesting eyes positioned on top of the head allowing to detect predators while being partially in the substrate. In contrast, arboreal species had great interocular and eye-nares distances, forming a more triangular head shape and reflecting their elongated and pointed head morphology.

This pattern aligns with observations of arboreal snakes, which often exhibit elongated heads that facilitate movement and navigation in complex three-dimensional environments (Jayne & Riley, 2007).

Overall, our results align with previous studies on morphological convergence in snakes. The elongated body proportions and slender morphology of arboreal species are consistent with findings on pythons, boas, and vipers, where elongation enhances climbing efficiency and maneuverability in complex three-dimensional environments (Pyron & Burbrink, 2009; Esquerré & Keogh, 2016; Alencar et al., 2017). Arboreal pythons and boas have independently evolved elongated bodies and reduced body mass to improve arboreal performance (Esquerré & Keogh, 2016), while vipers show morphological constraints that drive convergence in body shape despite phylogenetic differences (Alencar et al., 2017). The hydrodynamic adaptations of aquatic species align with findings on sea snakes where streamlined bodies, dorsally positioned nostrils, and laterally compressed tails optimize swimming (Brischoux & Shine, 2011). These adaptations improve propulsion and reduce drag, traits also observed in semi-aquatic species but to a lesser extent (Brischoux & Shine, 2011). Head morphology variation, including reduced inter-nares distances and specialized skull shapes, facilitate aquatic respiration and prey capture (Segall et al., 2016). The compact bodies of fossorial snakes support observations on burrowing species, which possess short, robust skulls and reinforced cranial structures to withstand mechanical stress (Strong et al., 2021). The wider bodies of burrowing species align with findings on fossorial and sand-dwelling reptiles, where a more cylindrical and compact body improves burrowing efficiency and reduces energy expenditure (Sharpe et al., 2015; Strong et al., 2021). The small body size of sand-dwelling species aligns with research on sand-swimming reptiles, where reduced size decreases friction with granular substrates and improves locomotion in loose sand (Sharpe et al., 2015). Slender, elongated bodies and smooth scales enhance movement efficiency by allowing sand-dwelling species to "swim" beneath the surface with minimal resistance (Sharpe et al., 2015).

Our results strengthen the idea that the physical constraints imposed by different habitats shape morphological disparity as evidenced by significant pairwise differences across most habitat groups (Table 5). However, two exceptions stand out: the lack of significant differences in disparity between terrestrial and semi-fossorial niches, and between fossorial and sand-dwelling niches. The first case may result from similar environmental pressures acting on the body, as both niches involve ground-dwelling lifestyles. The second could stem from the comparable biomechanical challenges imposed by sandy and subterranean habitats, where movement constraints and substrate interactions shape morphological adaptations in similar ways. These two same pairs correspond to those with the highest and lowest disparity scores respectively (Table 4; Figure 2). Thus, morphological disparity patterns reveal that semi-fossorial, semi-arboreal, and semi-aquatic species tend to resemble their terrestrial counterparts more than their fully specialized relatives. These observations support our hypothesis that highly specialized niches exhibit more homogeneous morphological features, whereas non-specialist habitat groups display significant morphological heterogeneity (Figure 3). Arboreal species exhibit an intermediate level of morphological disparity. However, among all ecological niches, arboreal species appear to exhibit the greatest gradient of diversity in morphological features. Similarly, aquatic species tend to exhibit a broader range of morphological features compared to semi-aquatic species. Sand-dwelling and fossorial ecotypes appear to be similarly disparate.

The test for morphological convergence in fossorial and aquatic species does not support a strong evolutionary convergence. This result suggests that while fossorial and aquatic environments impose significant biomechanical constraints on morphology resulting in reduced morphological disparity, species have evolved different solutions to these constraints rather than following a single convergent path. Semi-aquatic species also do not show a strong convergence. Similarly, semi-arboreal species show only moderate convergence, and their evolutionary timing (*P* = 0.281) does not support a strong historical trajectory toward a shared phenotype. Semi-aquatic and semi-arboreal species show intermediate levels of convergence, likely due to their mixed habitat use and a combination of ecological and functional constraints. The recent study of Tingle et al. (2024) aligns with our conclusion that functional diversity can lead to varied morphological outcomes rather than strict convergence. Furthermore, our findings align with studies which emphasize functional constraints over strict convergence, reinforcing the idea that biomechanical pressures do not always produce identical morphological adaptations (Vincent et al., 2006; Moon et al., 2019). However, recent studies highlighted strong convergence in head shape and feeding systems, suggesting that specific functional traits may be more prone to convergent evolution than overall morphology (Sherratt et al., 2018; Deepak et al., 2023).

Our findings provide a more nuanced understanding of the constraints imposed by specific ecological niches. While studies like those of Deepak et al. (2023) and Esquerré & Keogh (2016) focused on the convergence of head shape and body form in response to habitat use, our analysis extends these ideas by showing that morphological traits are not only convergent but also constrained by ecological factors. Our results add also a layer of complexity to the understanding of snake evolution, suggesting that while convergent evolution is common, the pathways leading to similar morphologies are often different.

Several studies on other taxa have also explored the relationship between morphology and ecological specialization and align with our findings. For example, fossorial mammals like moles and golden moles exhibit similar constraints on body shape and limb morphology due to burrowing demands, yet they achieve these adaptations through different evolutionary pathways (Samuels & Van Valkenburgh, 2009; Parker et al., 2013). Furthermore, research on anurans has demonstrated that arboreal species tend to exhibit more slender bodies and elongated limbs (Moen et al., 2016). These comparisons align with our findings that, across multiple taxa, ecological pressures drive morphological adaptations, but evolutionary history influences the specific solutions that arise.

While our study provides valuable insights into how habitat use drives morphological convergence in snakes, there are limitations in the study design and data analysis. One significant limitation is the exclusion of dietary factors from our analysis. Diet plays a critical role in shaping snake morphology, particularly head size and shape, as snakes adapt to different prey types and foraging modes (Herrel et al., 2008; Fabre et al., 2016; Sherratt et al., 2018; Deepak et al, 2023). By focusing primarily on habitat use, we may have overlooked how variation in diet contributes to morphological diversity. For example, snakes that specialize in eating large prey may develop broader heads, while those that feed on burrowing prey might evolve smaller heads. Incorporating dietary data could provide a more comprehensive understanding of the factors driving morphological convergence. Future research should integrate dietary information to explore how diet-specific adaptations influence morphological convergence, as an understanding the relationship between habitat use, prey type, foraging mode, and head morphology is essential for revealing how ecological pressures related to feeding drive morphological evolution. Additionally, while our study provides a detailed morphometric analysis, future research could benefit from more precise descriptions using geometric morphometrics to map morphological convergence with higher precision (Melchionna et al., 2021).

## Conclusion

Our study underscores the critical role of habitat use in shaping morphological convergence in snakes, revealing how similar ecological pressures can drive the evolution of comparable traits across distinct lineages. By examining variation in body and head shape across a diverse range of species, we provide insights into the complex interplay between morphology, ecology, and evolutionary history. While some ecological niches promote morphological homogeneity, others allow for greater disparity, reflecting differing degrees of functional constraints. Importantly, our findings suggest that convergence in snake morphology is not uniform but instead results from multiple adaptive pathways rather than a single evolutionary trajectory. This nuanced perspective highlights the importance of considering both ecological constraints and functional diversity when interpreting patterns of morphological evolution. Future research integrating dietary factors, geometric morphometrics, and finer-scale ecological data will further refine our understanding of how snakes—and other taxa—navigate the balance between specialization and morphological diversity in response to their environments.

## Supplementary data

**Table S1:** Taxa from the dataset with non-adult individuals.

| **Taxon** | **N** | **Life stage** |
| --- | --- | --- |
| *Boa constrictor* | 4 | Juvenile-young adult |
| *Elaphe quatuorlineata* | 2 | Young adult |
| *Eunectes murinus* | 1 | Juvenile |
| *Eunectes notaeus* | 1 | Juvenile |
| *Liasis olivaceus* | 2 | Young adult |
| *Malayopython reticulatus* | 3 | Young adult |
| *Pituophis melnaleucus* | 2 | Juvenile |
| *Python sebae* | 2 | Juvenile |

**Table S2:** Definitions of habitat groups.

| **Niche** | **Definition** |
| --- | --- |
| Terrestrial | Species that crawl on the ground and that rarely climb. |
| Sand-dwelling | Species that live in sandy environments such as deserts and dunes. |
| Semi-arboreal | Species that are terrestrial but with the ability to climb and move freely in vegetation. |
| Arboreal | Species that live mostly in trees and rarely come down to the ground. |
| Semi-aquatic | Species that depend on aquatic areas, mainly to feed on aquatic prey, but are generalist enough to leave these areas without difficulty. |
| Aquatic | Species that depend nearly exclusively on water. |
| Semi-fossorial | Species that live underground opportunistically or seek their food in an underground environment such as leaf litter, roots, rocks, etc... |
| Fossorial | Species that live exclusively underground. |

**Table S3:**
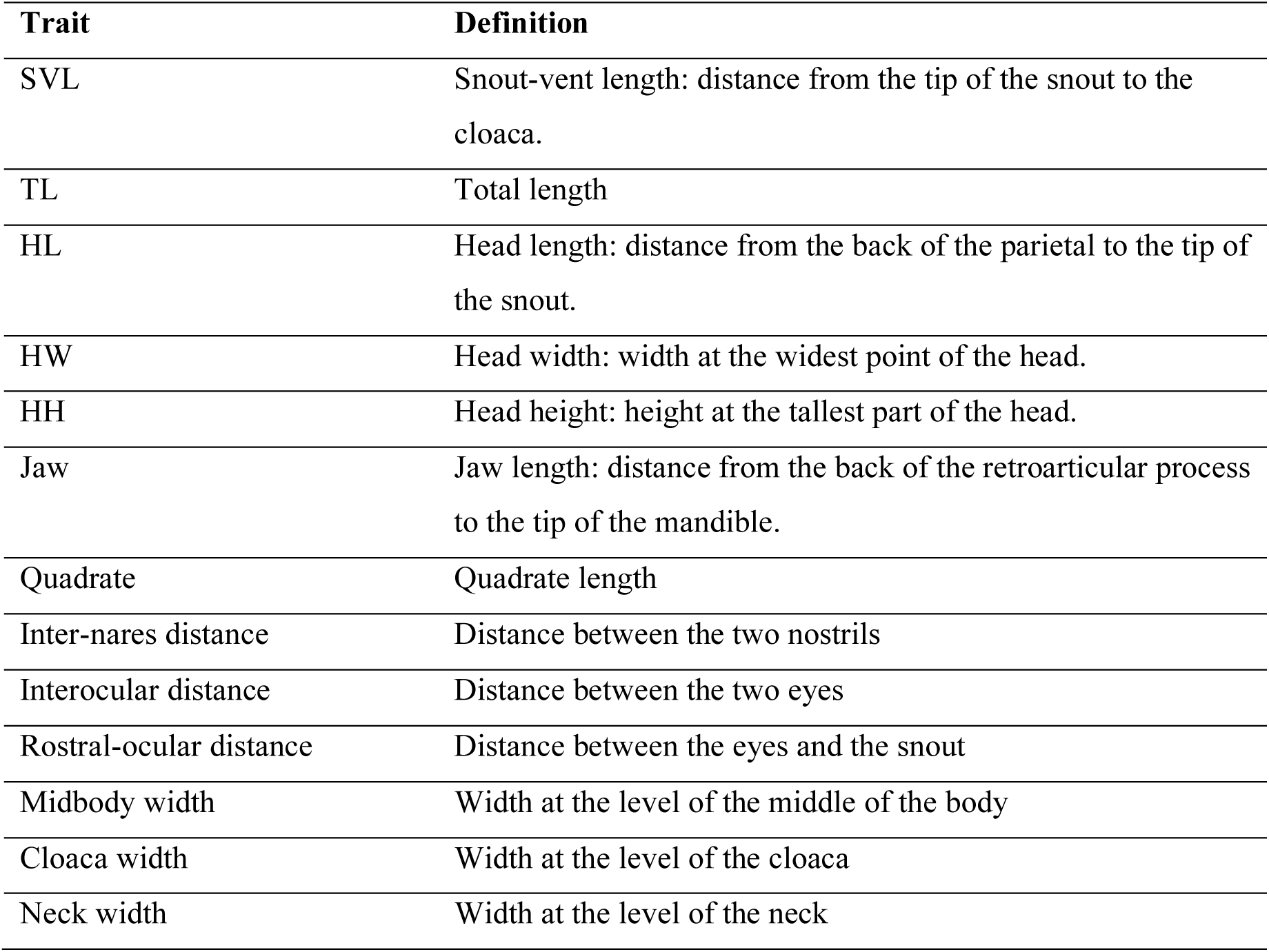
Morphological traits collected for morphometric analysis.

| **Trait** | **Definition** |
| --- | --- |
| SVL | Snout-vent length: distance from the tip of the snout to the cloaca. |
| TL | Total length |
| HL | Head length: distance from the back of the parietal to the tip of the snout. |
| HW | Head width: width at the widest point of the head. |
| HH | Head height: height at the tallest part of the head. |
| Jaw | Jaw length: distance from the back of the retroarticular process to the tip of the mandible. |
| Quadrate | Quadrate length |
| Inter-nares distance | Distance between the two nostrils |
| Interocular distance | Distance between the two eyes |
| Rostral-ocular distance | Distance between the eyes and the snout |
| Midbody width | Width at the level of the middle of the body |
| Cloaca width | Width at the level of the cloaca |
| Neck width | Width at the level of the neck |

**Figure S1:**
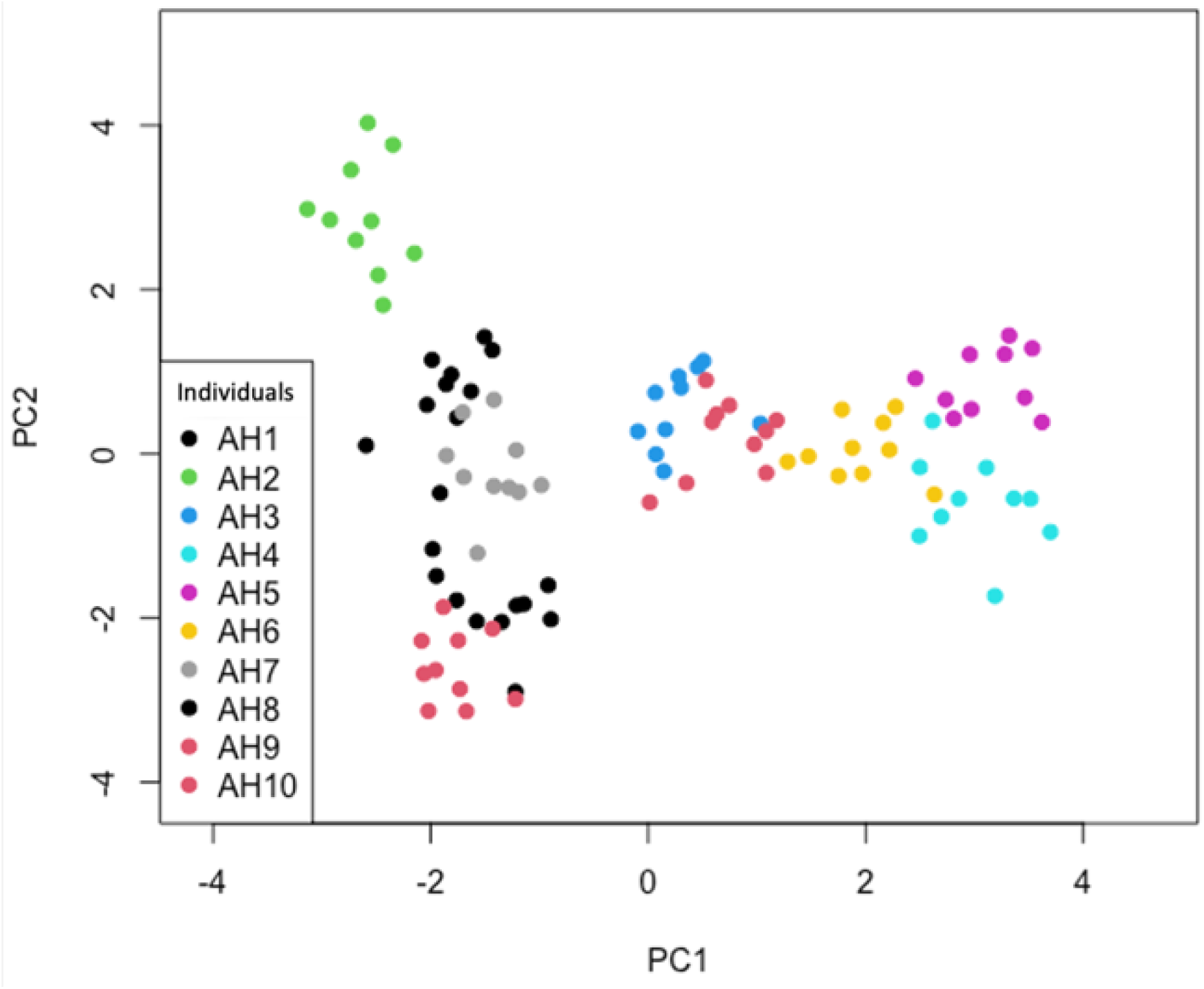
Repetition test results showing ten juvenile specimens of *Trimeresurus albolabris* on the PCA plot, indicating consistent morphological clustering with only minor overlap between individuals.

**Table S4:** Taxa from the dataset replaced by non-measured closely related taxa occurring in the phylogenetic tree.

| **Taxon** | **Replacement taxon** |
| --- | --- |
| *Atheris mongoensis* | *Atheris rungweensis* |
| *Atractaspis reticulata* | *Atractaspis aterrima* |
| *Atractus crassicaudatus* | *Atractus dunni* |
| *Grayia caesar* | *Grayia tholloni* |
| *Lamprophis littoralis* | *Boaedon radfordi* |
| *Limnophis bicolor* | *Natriciteres fuliginoides* |
| *Liophis taeniurus* | *Erythrolamprus juliae* |
| *Lycodon cardamomensis* | *Lycodon stormi* |
| *Macrocalamus schulzi* | *Oreocalamus hanitschi* |
| *Naja nana* | *Naja multifasciata* |
| *Philothamnus irregularis* | *Philothamnus angolensis* |
| *Toxicodryas adamanteus* | *Toxicodryas pulverulenta* |
| *Crotaphopeltis degeni* | *Crotaphopeltis tornieri* |

**Table S5:**
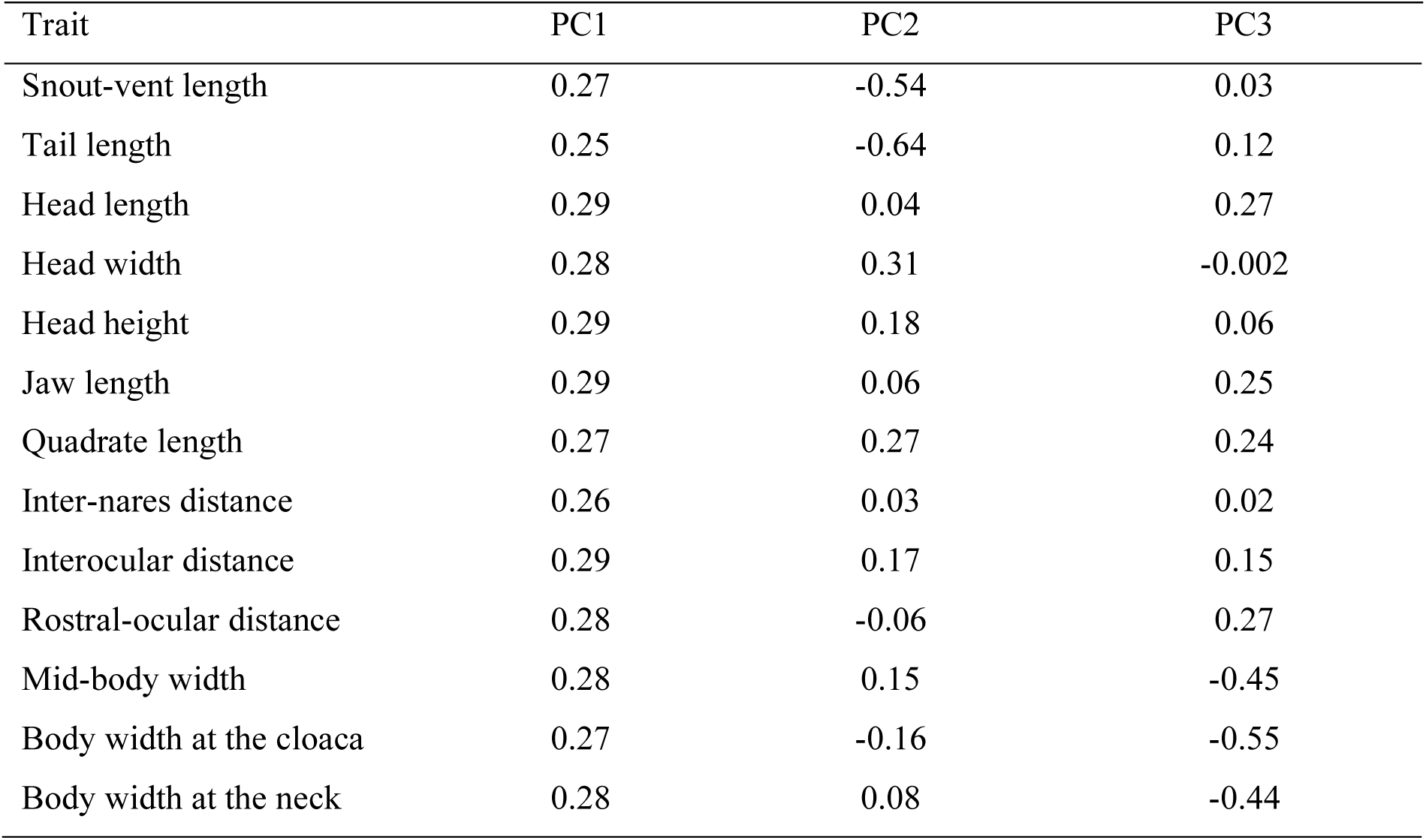
Loadings of Morphological Traits on the First Three Principal Components of Principal Component Analysis (PC1, PC2 and PC3)

**Table S6:**
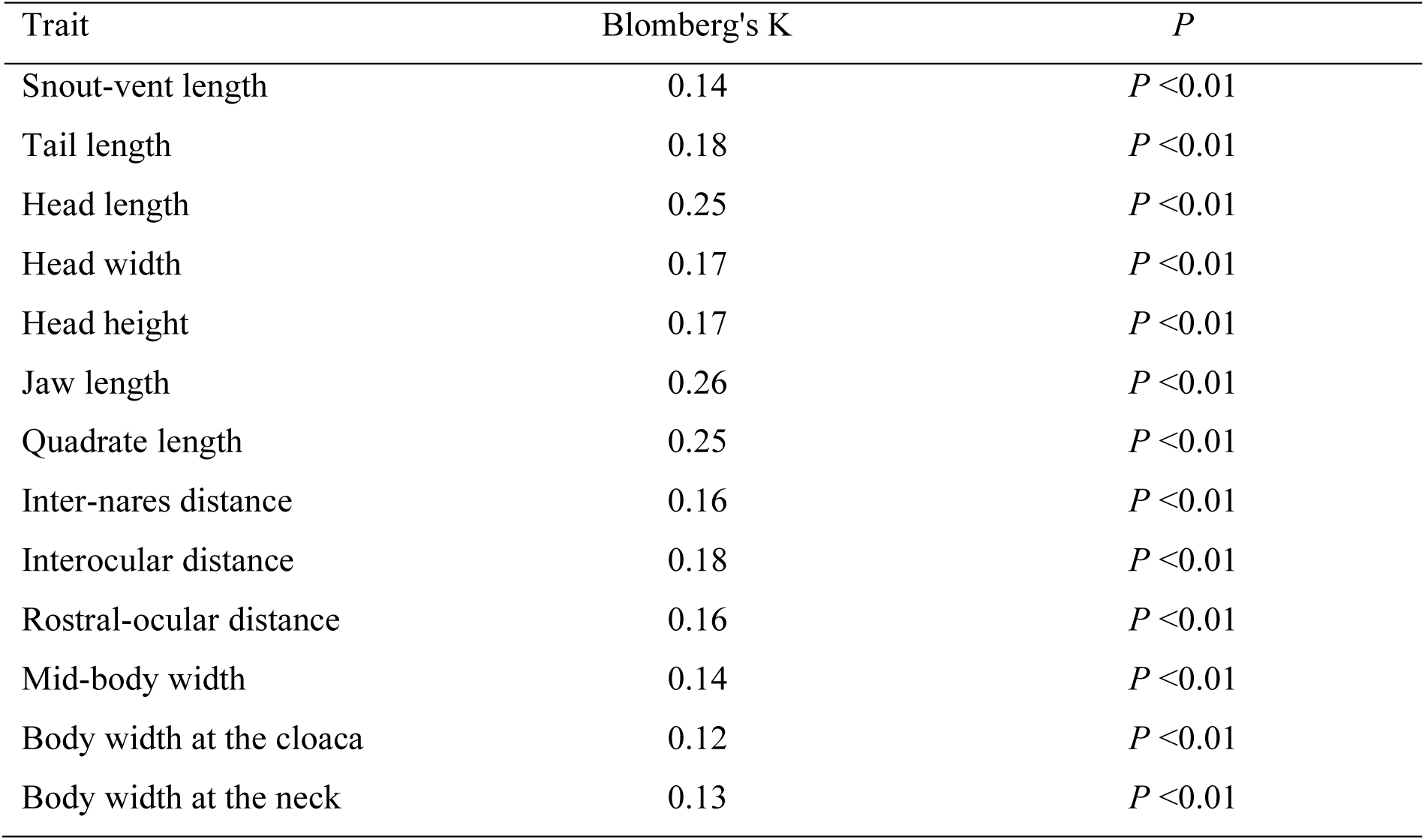
Blomberg’s K values and significance.

**Figure S2:**
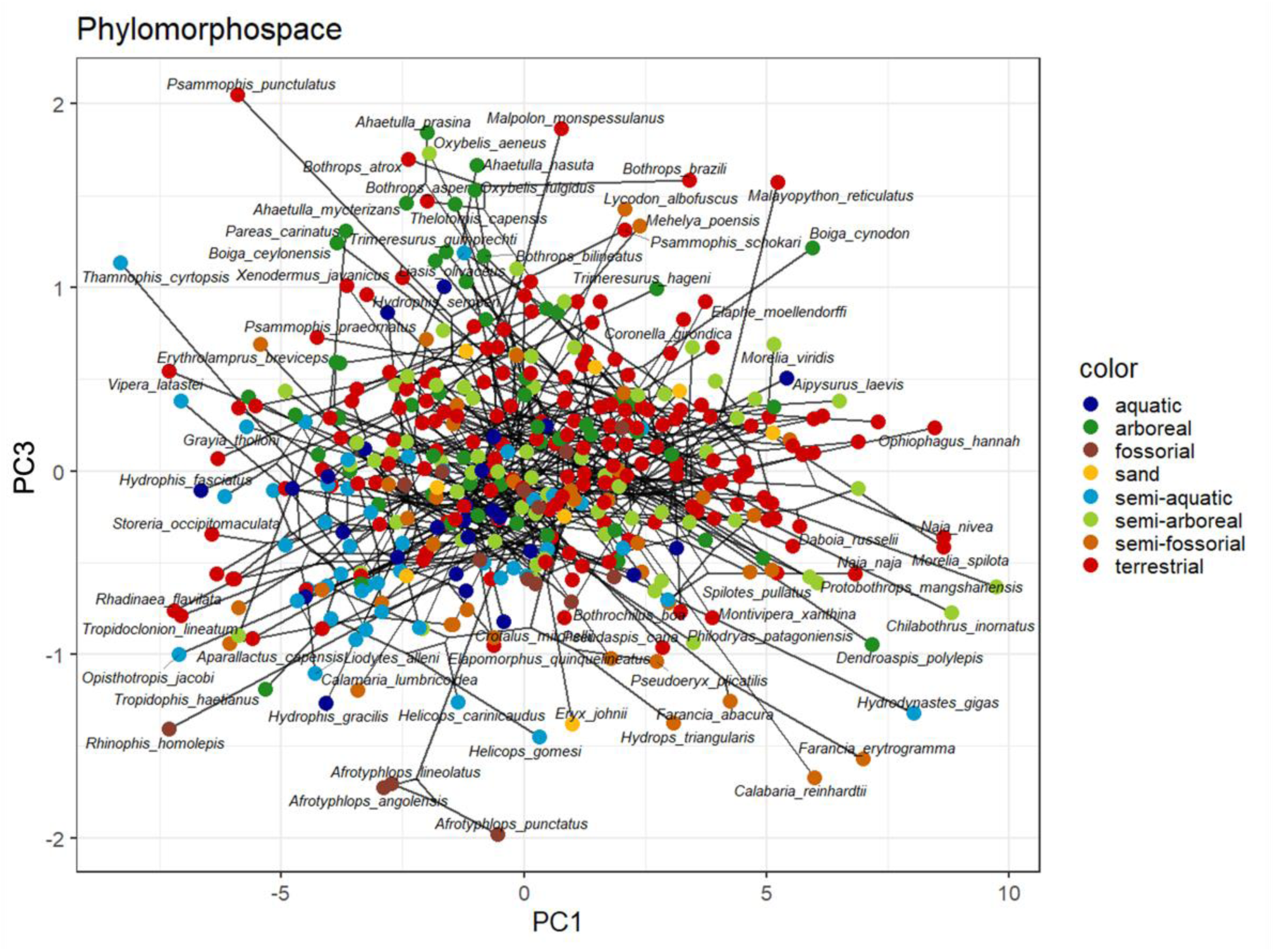
Phylomorphospace illustrating body and head shape variation across 436 species of snakes. The phylogeny is plotted in the morphospace described by axes 1 and 3. Nodes are colored based on symmetric rates ancestral-state estimation for habit mapped to the phylogeny.

**Figure S3:**
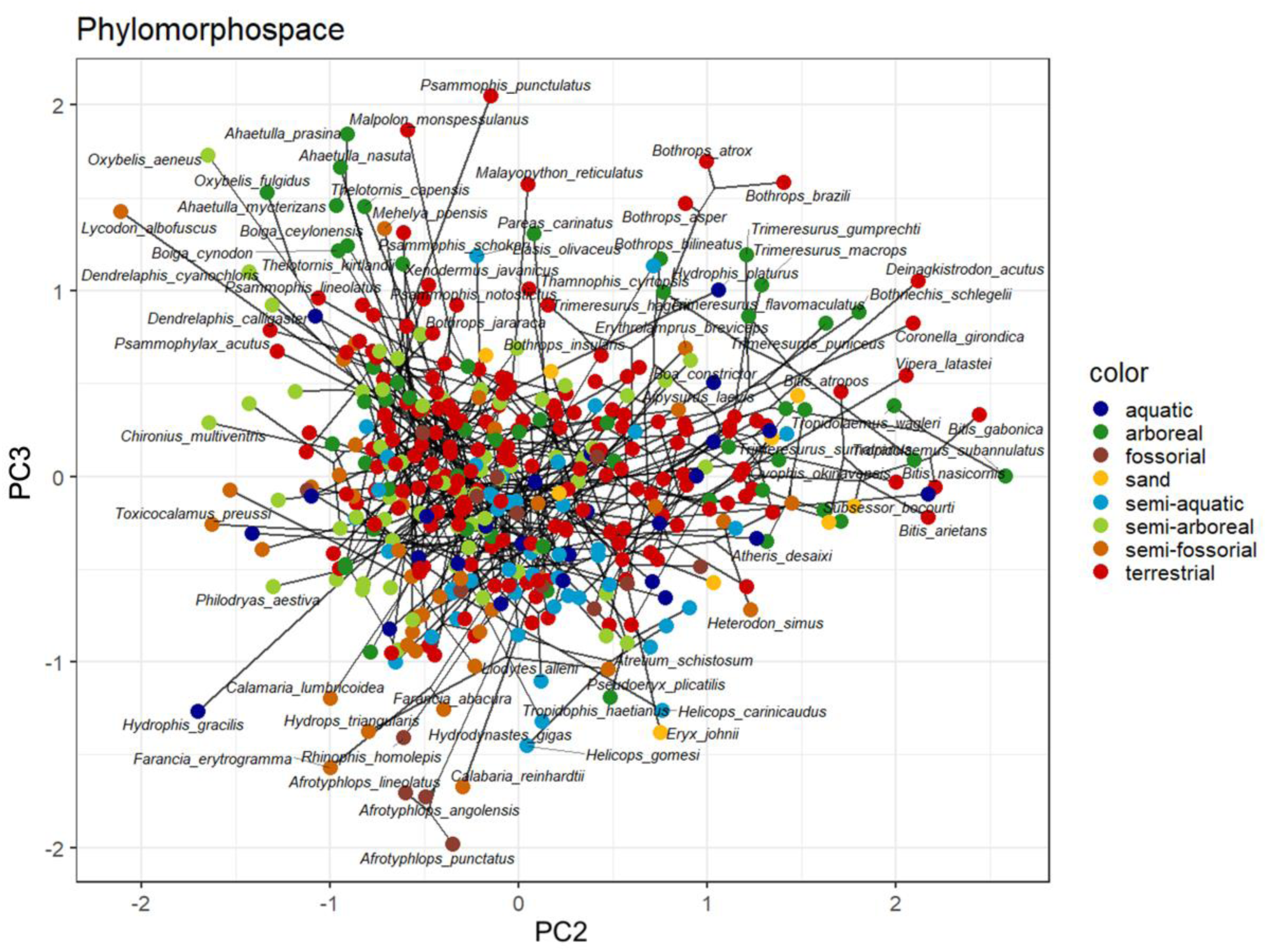
Phylomorphospace illustrating body and head shape variation across 436 species of snakes. The phylogeny is plotted in the morphospace described by the axes 2 and 3. Nodes are colored based on symmetric rates ancestral-state estimation for habit mapped to the phylogeny.

